# Leptomeningeal Neural Organoid (LMNO) Fusions as Models to Study Meninges-Brain Signaling

**DOI:** 10.1101/2023.12.01.569648

**Authors:** Hannah E Jones, Gabriella L Robertson, Alejandra Romero-Morales, Rebecca O’Rourke, Julie A Siegenthaler, Vivian Gama

## Abstract

Neural organoids derived from human induced pluripotent stem cells (iPSCs) provide a model to study the earliest stages of human brain development, including neurogenesis, neural differentiation, and synaptogenesis. However, neural organoids lack supportive tissues and some non-neural cell types that are key regulators of brain development. Neural organoids have instead been co-cultured with non-neural structures and cell types to promote their maturation and model interactions with neuronal cells. One structure that does not form *de novo* with neural organoids is the meninges, a tri-layered structure that surrounds the CNS and secretes key signaling molecules required for mammalian brain development. Most studies of meninges-brain signaling have been performed in mice or using two-dimensional (2D) cultures of human cells, the latter not recapitulating the architecture and cellular diversity of the tissue. To overcome this, we developed a co-culture system of neural organoids generated from human iPSCs fused with fetal leptomeninges from mice with fluorescently labeled meninges (*Col1a1-GFP*). These proof-of-concept studies test the stability of the different cell types in the leptomeninges (fibroblast and macrophage) and the fused brain organoid (progenitor and neuron), as well as the interface between the organoid and meningeal tissue. We test the longevity of the fusion pieces after 30 days and 60 days in culture, describe best practices for preparing the meninges sample prior to fusion, and examine the feasibility of single or multiple meninges pieces fused to a single organoid. We discuss potential uses of the current version of the LMNO fusion model and opportunities to improve the system.

## Introduction

The development of the human central nervous system (CNS) is a complex process requiring the precise coordination of signaling molecules and various cell types. Our understanding of human CNS development is mostly derived from rodent and non-human primate models, along with investigation of post-mortem human brain tissue. While informative, studies using these approaches do not adequately mimic the *in vivo* processes of human brain development. Thus, neural organoids derived from human pluripotent stem cells, both embryonic stem cells (ESCs) and iPSCs, have become useful tools for understanding various stages of human brain development. Neural organoids can be used to model aspects of human CNS development, such as neurogenesis and synaptogenesis, and produce a heterogenous mixture of cell types^1–3^. Neural organoids can form distinct layers reminiscent of cortical layers found *in vivo* and bear an organizational and transcriptomic resemblance to the gestational human cortex^3–5^. Additionally, small molecules and other morphogens can be used to pattern the organoids into specific cell fates or encourage the patterning into different developmental regions, such as telencephalic or cerebellar-like neural organoids^2,6,7^.Thus, neural organoids have become central tools for studying early human CNS developmental processes and for translational applications.

One major drawback of current human neural organoid platforms is their lack of support structures that are required for proper CNS development and function *in vivo*^2,8,9^. These are structures or cell types that are not derived from the neuroectoderm that give rise to the CNS and include vasculature, microglia, and meninges. Some strides have been made to incorporate microglia and vasculature into neural organoids^9,10^ While these systems have improved our understanding of interactions between neural and non-neuronal cell types within neural organoid models of human CNS development, these platforms still lack many of the characteristics and spatial organization of human brain morphology. A structure that is key to proper formation and maintenance of the CNS *in vivo* is the meninges, a tri-layer structure composed of fibroblasts, blood vessels, and immune cells that encases the entire CNS. It has been well established that the meninges play an essential role in brain development by secretion of important factors: meningeal retinoic acid is required for cortical and cerebellar development in mice and humans^11^, meningeal CXCL12 directs Cajal-Retzius cell migration^12^, and the pia layer of the meninges forms the basement membrane interface which is required for radial glial cell attachment^13^. Despite this, few attempts have been made to incorporate meningeal tissue into neural organoid models and much of the work to understand interactions between the meninges and neuronal cell types during development have been conducted in rodent models or meningeal cell lines.

Overall, there is a need for human neural organoid systems to better recapitulate the structure and organization of the human brain during development, and we propose the meninges as a key component to achieve this complexity. Here, we describe a three-dimensional (3D) co-culture system created from iPSC-derived human neural organoids fused with embryonic leptomeninges from *Col1a1-GFP* mice, which we named Leptomeningeal Neural Organoid (LMNO) fusions. We show that the meningeal compartment of LMNOs retains key characteristics of leptomeninges *in vivo*, such as layer-specific fibroblast markers and resident immune cells, up to 60 days in culture. We also find that neurons in the LMNO fusions undergo less cell death compared to neurons in non-fused organoids. Finally, we find evidence of iPSC-derived human brain organoid Cajal-Retzius-like neuronal cells migrating into the meningeal compartment of LMNOs, potentially responding to Cxcl12 ligand derived from meningeal cells. This proof-of-concept study provides an important initial framework for incorporation of meninges into brain organoid platforms.

## RESULTS

### Cellular composition of neural organoids and leptomeningeal tissue

Prior to fusing, day 15 human neural organoids have begun to develop ventricular-like zones, which recapitulate the developing cytoarchitecture of the human cortex (**Fig. 1A**). The cells surrounding these ventricular-like zones express the expected markers of neural progenitors such as PAX6 and SOX2 (**Fig. 1A**). By day 15, MAP2^+^ neurons have started to differentiate and migrate away from the ventricular zone (**Fig. 1A**). This organization in neural organoids is intrinsic and develops prior to our fusion; however, minimal neurogenesis has occurred at this timepoint.

**Figure 1:**
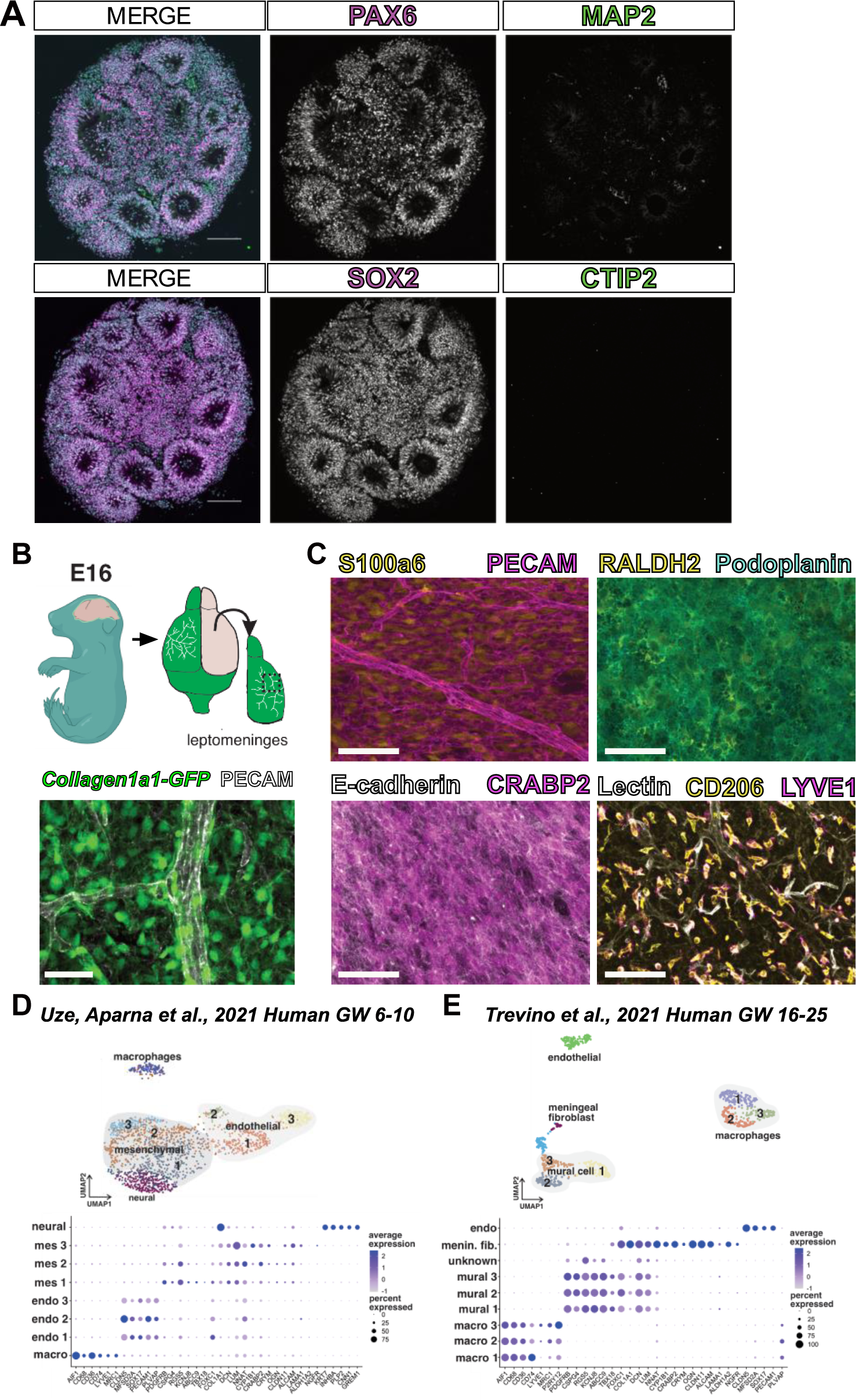
Characterization of neural organoids and mouse fetal meninges prior to fusion. (**A**) Representative images of day 15 neural organoid labeled with PAX6/MAP2 or SOX2/CTIP2, scale bars 100 μm. (**B**) Schematic and maximum-projection image depicting isolation and whole-mounting of mouse embryonic *Col1a1-GFP+* leptomeninges, image shows GFP+ leptomeningeal fibroblasts (green) and vasculature labeled with PECAM (white). (**C**) Identification of cell types in mouse leptomeningeal whole-mount tissue, images show labeling of S100a6 (fibroblasts)/PECAM (vasculature), RALDH2 (fibroblasts)/Podoplanin (fibroblasts), E-cadherin (arachnoid barrier)/CRABP2 (fibroblasts), and CD206 (macrophages)/LYVE1 (macrophages)/Lectin (vasculature); all scale bars 100μm. (**D-E**) Dot-plots showing conservation of mouse meningeal cell types and markers in human brain data sets at (**D**) GW 6-10 (from Uze, Aparna et. al. 2021) and (**E**) GW 16-25 (from Trevino et. al. 2021).

The leptomeninges is comprised of the inner two layers of the meninges, the pia and arachnoid. It contains the pial and arachnoid fibroblasts that produce ECM, retinoic acid, CXCL12, and BMPs vital for brain development. The mouse leptomeninges can be isolated as a separate structure from the brain and the outer layer of the meninges, the dura. We isolated meninges from the *Collagen1a1-GFP (Col1a1-GFP)* reporter mouse line in which all fibroblasts are labeled with GFP (**Fig. 1B, schematic**); using previously described whole-mounting techniques^14^ combined with immunofluorescent labeling we can visualize *Col1a1-GFP+* fibroblasts and the associated meningeal vasculature (**Fig. 1B**). Single-cell profiling studies of the meninges reveal unique layer-specific fibroblast subtypes and tissue-resident macrophages called border associated macrophages (BAMs). These cell populations can be visualized in the dissected leptomeninges using specific markers: pia-layer fibroblasts labeled by S100a6, arachnoid fibroblasts labeled with RALDH2 and PDPN, pseudo-epithelial arachnoid barrier cells labeled by ECAD and CRABP2, and BAMs labeled by CD206 and LYVE1 (**Fig. 1C**).

We previously showed that some meningeal fibroblast layer specific markers (e.g., S100a6, CRABP2) are shared between fetal mouse and human meninges^15^. To further test this, we used published single nuclear RNA sequencing data sets from early embryonic (human gestational week 6-10) and later fetal (gestational week 16-25) human brain development to identify cell clusters that are representative of putative meningeal mesenchymal/fibroblast, meningeal macrophage, and vascular cells (**Fig. 1D, E**). At early embryonic stages, three mesenchymal cell clusters were identified with one of the clusters (mes 3) showing enriched gene expression of previously identified mouse fetal pia (*NGFR, LAMA1, CYP1B1, LUM*) and arachnoid cells (*CLDN11, ALCAM*)^15^ (**Fig. 1D**). The analysis of clusters at later stages of fetal human brain development identified a meningeal fibroblast enriched cluster, likely representing a mixture of arachnoid and pial cells (**Fig. 1E**) as first identified in mouse fetal meninges. At both stages, we identified endothelial cell clusters with CNS specific gene expression profiles, (low *PLVAP*, high *CLDN5, SOX17*) and macrophage clusters with *MRC1* and *LYVE1* (**Fig. 1D, E**), consistent with studies in mouse on vascular and macrophage development. This indicates that mouse leptomeninges contain meningeal fibroblasts, macrophages, and vasculature populations that are transcriptionally like human embryonic and fetal meninges. Further, these data provide supportive evidence that mouse leptomeninges are a reasonable proxy for human-derived tissue.

### Generation of LMNO fusions

Neural organoid platforms mimicking human development have yet to incorporate the meninges, a structure essential for proper brain development. To develop LMNO fusions, neural organoids were first generated from iPSCs as previously described^3,16^ and grown up to day 15 in culture, resembling approximately 6-9 gestational weeks in human (**Fig. 2A**). Then, we isolated leptomeninges from embryonic day (E)16 *Col1a1-GFP+* mice, in which meningeal fibroblasts express GFP under the *Col1a1* promoter^17^. Prior to isolation of the meninges, embryos were perfused with ion-free HBSS on ice to remove blood contents (**Fig. 2A**). Then, brains were removed and the leptomeninges were dissected from each hemisphere of the forebrain (**Fig. 2A**), as previously described^14^. Isolated meninges were collected in sterile, ion-free HBSS on ice until ready to be transferred into culture with neural organoids (**Fig. 2A**). We transferred one neural organoid along with 2-4 pieces of *Col1a1-GFP+* forebrain leptomeninges into one well of a static culture system consisting of a round-bottom Eppendorf tube secured with hot glue to the bottom of each well of a 6-well plate (**Fig. 2A**). Each well was filled with approximately 300-500uL of organoid differentiation media and given complete media changes every other day (**Fig. 2A**). We monitored neural organoid-meninges fusion progression by capturing bright-field images every other day (**Fig. 2A-B**). At day 3 in culture, the organoid and meninges components remain mostly separate, but by day 6 we observe fusion between the neural organoid and meninges, thus generating LMNOs (**Fig. 2B**). LMNOs remained fused throughout the course of the experiment, including after collection for staining at day 30 or 60 (**Fig. 2A-B**).

**Figure 2:**
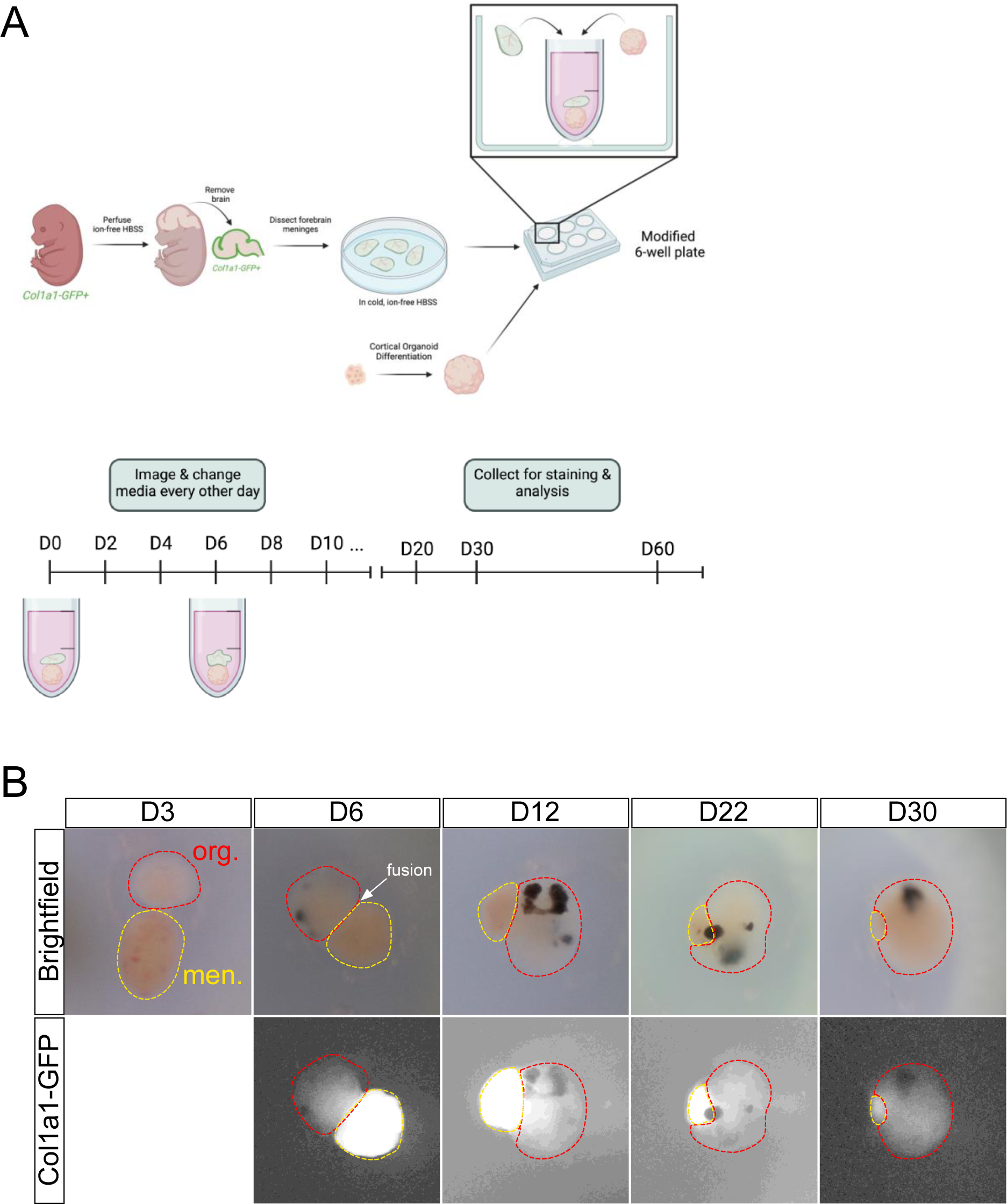
Leptomeningeal-neural organoid (LMNO) fusion culture establishment and growth. (**A**) Graphical depiction of leptomeningeal dissection and establishment of LMNO fusions cultures. (**B**) Representative stereoscope images of LMNO fusions throughout time in culture. Red-dashed lines highlight organoid component, yellow-dashed lines and *Col1a1-GFP* fluorescence distinguishes meningeal component.

### Characterization of specialized meningeal cell types within LMNO fusions

The meninges are a complex tissue consisting of a diverse array of cell types including specialized fibroblasts and tissue resident immune cells. Thus, we next sought to understand the cellular composition of the meningeal tissue compartment of the LMNO fusions.

The meninges contain tissue-resident yolk-sac derived macrophages called border associated macrophages (BAMs) that play an important role in immunological surveillance and response to infection and injury^18,19^. BAMs appear in the meninges embryonically persisting through adulthood and are identified by the markers CD206 (also known as mannose receptor C-type 1 (MRC1)) and lymphatic vessel endothelial hyaluronan receptor 1 (LYVE1). We conducted fluorescent immunolabeling for CD206 and LYVE1 on sections from LMNO fusions at day 30 and day 60 and found an abundance of CD206 and LYVE1 labeling of BAMs in the meningeal compartment of the LMNOs at both timepoints (**Fig. 3A**). The CD206 and LYVE1 labeling remained within the meningeal compartment and no labeling was observed in the organoid compartment, suggesting that any BAMs that persist in the LMNO fusions stay confined in the meninges. At day 60, we observed that much of the *Col1a1-GFP+* signal colocalized with CD206 and/or LYVE1 (**Fig. 3A**), suggesting that BAMs may be phagocytosing cells within the LMNOs.

**Figure 3:**
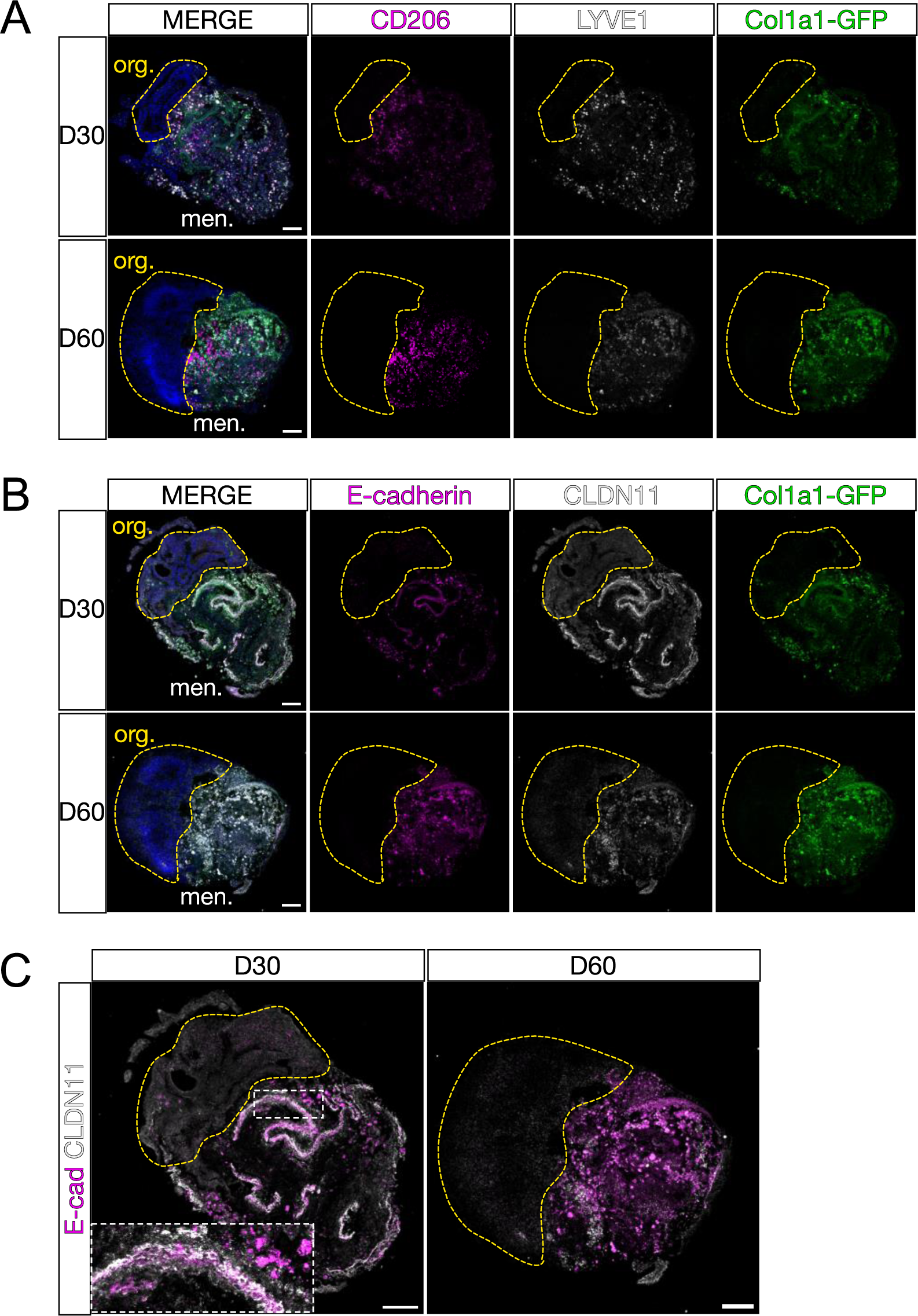
Characterization of meningeal components in LMNOs after 30 and 60 days in culture. Images showing LMNOs after 30 or 60 days in culture, labeled with markers for meningeal cell types; (**A**) macrophages labeled with CD206 (magenta) and LYVE1 (white), (**B**) arachnoid barrier cells labeled with E-cadherin (magenta) and CLDN11 (white); all fibroblasts labeled with *Col1a1-GFP* (green). (**C**) Images from **B** with E-cadherin/CLDN11 overlaid; inset shows overlap between markers. Yellow-dashed lines distinguish organoid compartment (org.) from meninges (men.). All scale bars 100μm.

The meninges also contain specialized fibroblasts called arachnoid barrier (AB) cells that are critical for maintaining the barrier properties of the meninges^20^. AB cells are specified and form elaborate cell-cell adhesions utilizing the molecules E-cadherin (ECAD) and claudin 11 (CLDN11) during embryonic development^20^. We conducted fluorescent immunolabeling for AB cell junctional proteins ECAD and CLDN11 on sections from LMNO fusions at day 30 and day 60 and saw labeling for these markers within the meningeal compartment at both timepoints (**Fig. 3B-C**). At day 30, we observed ‘ribbons’ of co-localized ECAD/CLDN11 labeling reminiscent of the AB layer *in vivo* (**Fig. 3B**, **Fig. 3C inset**). By day 60, these structures were mostly absent and most of the ECAD/CLDN11 labeling was seen in ameboid-like structures (**Fig. 3B-C**); given the abundance of BAMs we suspect that any AB structures that remain in the meningeal compartment of the LMNO fusions are phagocytosed by this timepoint. Taken together, this data shows that specialized meningeal cell types (BAMs and AB cells) are able to persist in LMNO fusion cultures at day 30 and 60 and retain functional and structural characteristics.

### Neural organoids in the LMNO fusion model display reduced apoptosis

LMNO fusions retain characteristic meningeal cell types (*Col1a1-GFP+* fibroblasts, AB cells, and BAMs) 30-60 days into culture, though we observe a significant amount of cell fragmentation and what appears to be phagocytosed cellular components by resident BAMs by day 60. Thus, we next quantified the relative frequency of cell death as a readout for overall health of the culture system. To do so, we conducted immunofluorescent labeling for the apoptosis pathway component activated Caspase-3, along with CRABP2, which labels both meningeal fibroblasts and neurons, in LMNO fusions at D30 and D60 (**Fig. 4A, LMNO**). We also conducted the same staining in neural organoids not fused to meninges but kept in the same culture conditions as the LMNO fusions (**Fig. 4A, org. only**). We then quantified cell death by measuring the percentage of the area labeled with Caspase-3 of each LMNO compartment (organoid, meninges) (**Fig. 4B**). At D30, we find that the organoid compartment of LMNOs contain significantly less Caspase-3 labeling compared to the meninges compartment, and the organoid compartment had significantly less Caspase-3 labeling compared to organoids lacking meninges (**Fig. 4B**). This effect was only observed at D30; at D60 we find that the amount of Caspase-3 labeling is not significantly different between the organoid compartment of LMNO fusions and organoids alone (**Fig. 4B**). From these results, we speculate the meninges component of LMNOs are protective against cell death within the organoid compartment, potentially via secretion of pro-survival signals.

**Figure 4:**
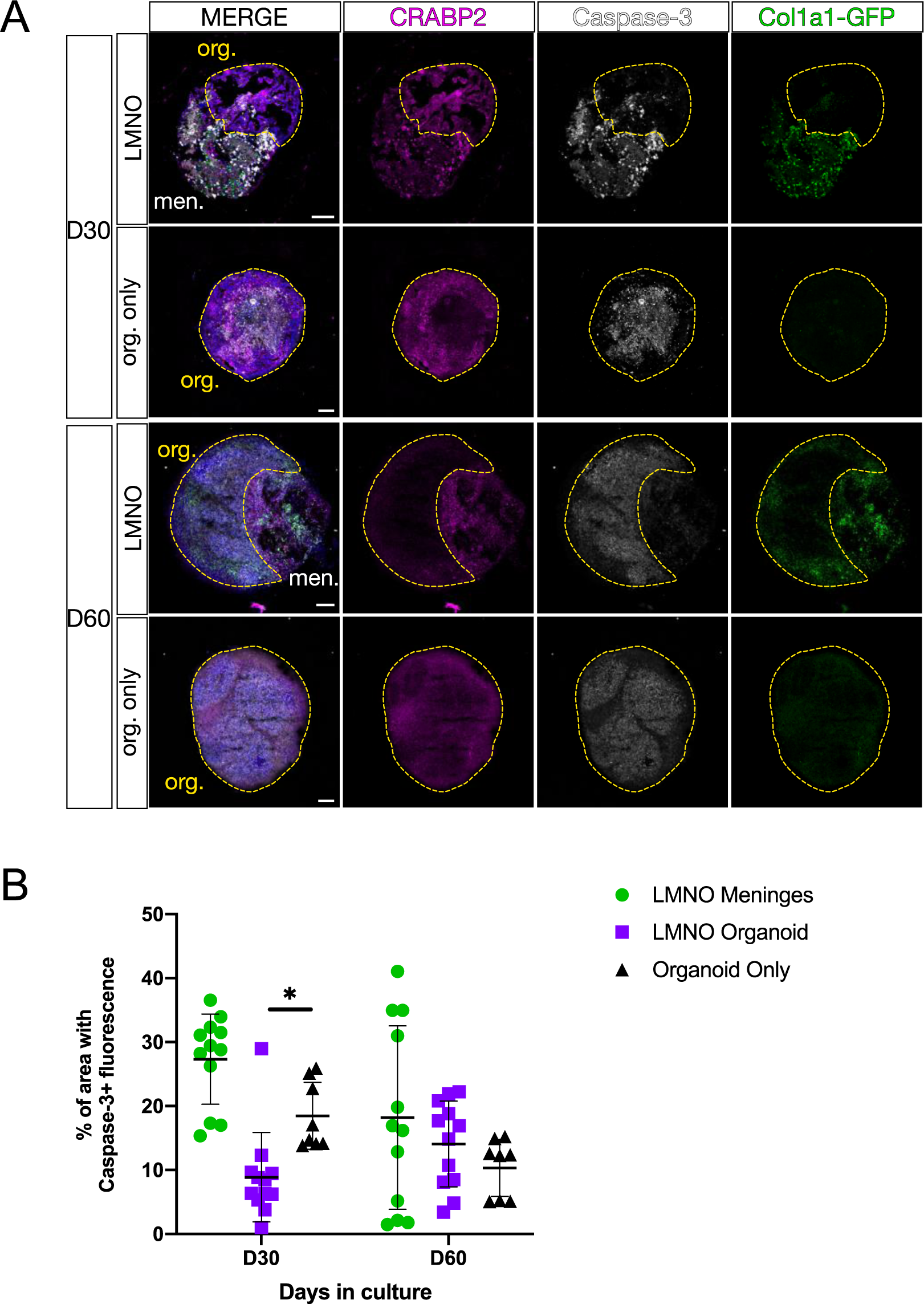
Neural organoids in LMNO fusions show reduced levels of apoptosis. **(A)** Images showing Caspase-3 (white) and CRABP2 (magenta) staining in LMNO fusions and non-fused organoids (org. only) at 30 and 60 days in culture, *Col1a1-GFP* (green) labeling fibroblasts. Yellow-dashed lines outline organoid compartment. Yellow-dashed lines distinguish organoid compartment (org.) from meninges (men.). All scale bars 100μm. (**B**) Quantification of percentage of area of organoid and meningeal compartments with Caspase-3 fluorescence at 30 and 60 days in culture; meninges in the LMNO fusion (green circles), LMNO fusions (purple squares), non-fused organoid (black triangle). Two-way ANOVA with multiple comparisons revealed significant differences between LMNO fusion versus organoid only at D30 (p = 0.0423) (p< 0.05, *).

### Human REELIN+ neurons migrate into meningeal tissue in LMNO fusion model

Given the evidence for neuronal cell migration in other iPSC-derived organoid models^21^, we examined whether any human cells migrated into the meningeal compartment of the LMNO fusion using a marker specific for human mitochondria. Human mitochondria were observed in the mouse meningeal tissue in some instances (**Fig. 5A-B**). These migratory human cells express REELIN (**Fig. 5C**), a marker of Cajal-Retzius-like cells known to be present in iPSC-derived neural organoids and to migrate within the developing human brain. These data suggest that there is integration of the two tissues and that human neural cells migrate into the meningeal tissue. Conversely, PAX6+ neural progenitors remain in the neural organoid around ventricular-like zones (**Fig. 5A**).

**Figure 5:**
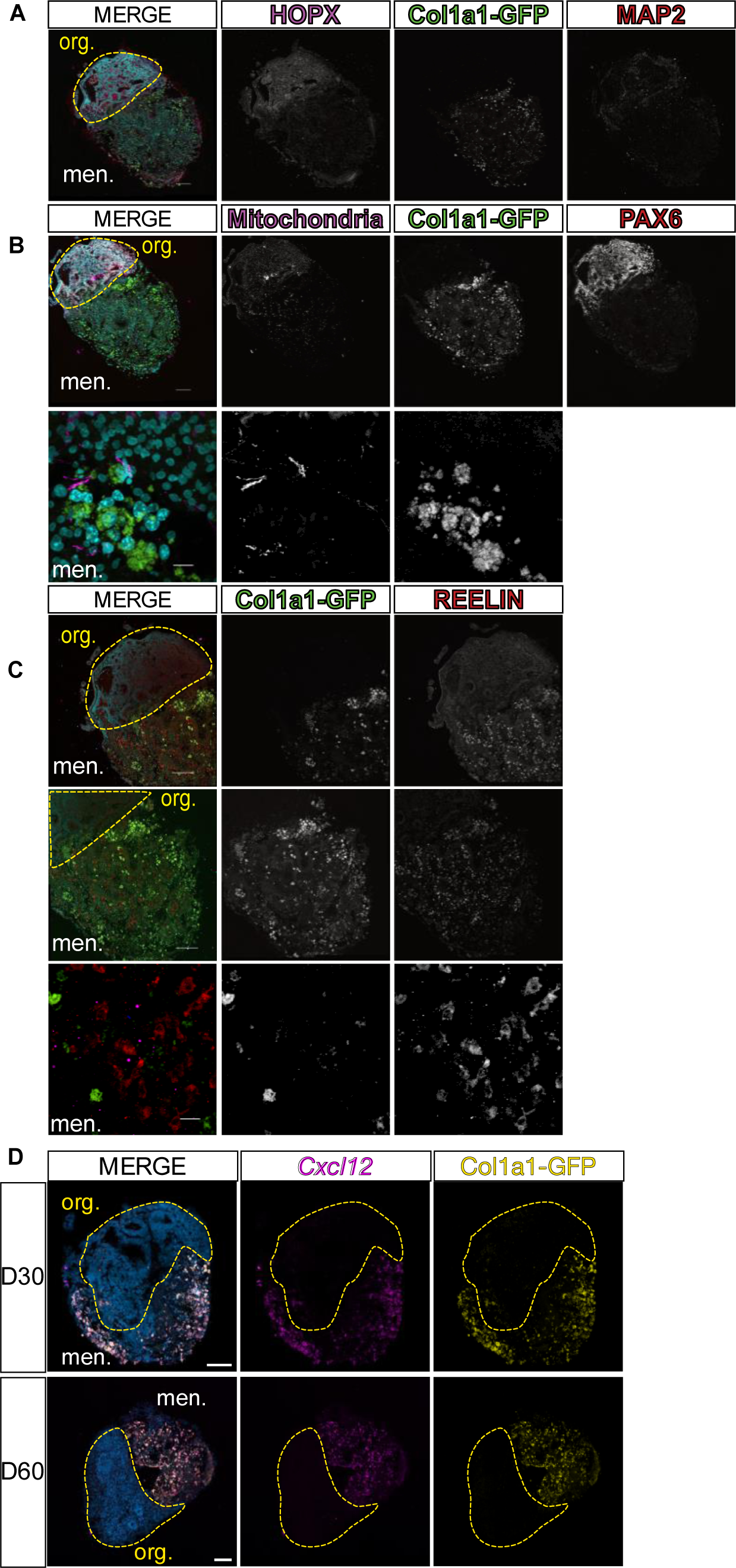
LMNO fusions recapitulate *in vivo* Cxcl12-dependent REELIN+ Cajal-Retzius cell migration. (**A-B**) Images of LMNO fusions labeled with (**A**) neuronal markers HOPX (magenta)/MAP2 (red), **(B)** progenitor marker PAX6 (red) and human mitochondria (magenta). Top image scale bar 100μm and bottom image scale bar 10μm. (**C**) Cajal-Retzius cell marker REELIN (red); *Col1a1-GFP* marking meningeal fibroblasts (green). Top 2 images scale bar 100μm and bottom image scale bar 10μm. (**D**) Images of *in situ* detection of *Cxcl12* (magenta) in LMNO fusions at 30 and 60 days in culture, *Col1a1-GFP* marking meningeal fibroblasts (yellow). Yellow-dashed lines distinguish organoid compartment (org.) from meninges (men.). All scale bars 100μm.

It is well established that meningeal-derived factors are essential for proper cortical development. One essential factor, CXCL12, is secreted by the meningeal fibroblasts embryonically and is required for Cajal-Retzius cell migration *in vivo*^12^. Given that we observe organoid-derived REELIN+ Cajal-Retzius-like cells within the meningeal compartment of LMNOs, we next sought to determine if the LMNO meningeal compartment acts as a source for CXCL12 ligand. To do so, we conducted RNAscope *in situ* hybridization for *Cxcl12* mRNA on LMNO fusions at D30 and D60. At both timepoints, we observe abundant *Cxcl12* signal within the meningeal compartment of LMNOs, and signal is notably lower in the organoid compartment (**Fig. 5C**). This may suggest a CXCL12-dependent mechanism is driving Reelin+ cell migration into the meningeal compartment of LMNO fusions.

## Discussion

We present a method for co-culturing human iPSC-derived neural organoids with embryonic mouse meninges, referred to as LMNO fusions. In this system, embryonic mouse leptomeninges placed in culture with neural organoids undergo fusion in six days and can remain in culture through 60 days. We find that the leptomeningeal compartment of LMNO fusions retain key characteristics of the meninges *in vivo*, with tissue-resident macrophages and specialized arachnoid barrier cells persisting in culture through 30 days. Furthermore, the cell types known to be present in neural organoids were present after fusion, suggesting that fusion did not disrupt neural organoid development. We also find that the leptomeninges are potentially neuroprotective in the LMNO organoid compartment within the first 30 days in culture. Finally, we show that the LMNO organoid compartment recapitulates characteristics of cortical development *in vivo*, including producing migratory REELIN+ Cajal-Retzius-like cells that potentially respond to LMNO meninges compartment-derived CXCL12.

Our method provides a novel framework for the co-culturing of neural organoids with leptomeningeal tissue. In the LMNO system, whole meninges are placed into culture with neural organoids and undergo spontaneous fusion. Although the meninges in LMNOs do retain key cell types (**Fig. 3**), they do not retain their overall organization or form key structures at the interface between the meningeal and organoid components. *In vivo*, the meninges consist of three layers— the pia, arachnoid, and dura—with the pia layer residing closest to the brain surface, forming the glia limitans and acting as a crucial attachment point for radial glial cells^13^. Future meninges-neural organoid platforms should aim to retain this organization, via scaffolding and/or methods to self-organize the meninges into the proper layer order around organoids. A dissociated meningeal cell line has been used to coat neural organoids with meningeal fibroblasts^22^; future systems could employ similar methods with primary embryonic meningeal cells to maintain the cellular diversity.

Human leptomeningeal tissue has yet to be derived from iPSCs *in vitro*. However, the LMNO fusion model allows us to study the interaction of human neural organoids and mouse leptomeningeal tissue *in vitro*, which may give insight into key signaling pathways that are essential to deriving these tissues. Furthermore, this model provides evidence that meningeal tissue can be cultured *in vitro* and retain the cellular diversity of the tissue. These future, improved systems will not only be important for modeling brain development *in vitro* but will also be useful models for studying meningeal development and brain-meninges cross-talk.

Within the meningeal component of LMNO fusions, we observed the survival of meningeal tissue-resident macrophages, BAMs, marked by expression of CD206 and LYVE1. The BAMs appeared to remain confined to the meninges and were not observed in the organoid compartment of LMNO fusions, mirroring the homeostatic localization of BAMs *in vivo.* We also observed GFP+ fluorescence within the CD206+/LYVE1+ cells, suggesting that the macrophages present in the LMNO fusions may actively phagocytose fragments of dead fibroblasts. We also detect cells expressing the junctional proteins E-cadherin and Claudin-11, which are expressed by AB cells *in vivo.* These AB-like structures are seen at day 30 but appear to break down by day 60 in culture. It has been established that the AB layer begins establishing cell-cell junctions along with its barrier function embryonically, however it is unknown what signals contribute to maintenance of the AB layer *in vivo.* Potentially, other meningeal fibroblasts present in the LMNO meninges compartment produce pro-survival signals or the BAMs act to reduce cytotoxicity by phagocytosing dead cells.

During development, the meninges secrete factors (CXCL12, retinoic acid, BMPs) that are essential for proper cortical development. In the LMNO fusion system, we observed that the meninges component produces CXCL12 ligand, which *in vivo* is necessary for driving Cajal-Retzius cell migration. We also found evidence that the meninges promote cell survival in the organoid of the LMNO fusion through day 30 in culture. Potentially, other meningeal-derived cues promote neuronal survival in LMNO fusions, though these mechanisms do not seem to persist through day 60 in culture given the widespread cell death seen at this timepoint.

Developmental apoptosis is an evolutionary conserved pathway critical for neurodevelopment^23^, that is actively regulated during the generation of neural organoids^24^. However, increased cell death can also arise from long-term culturing conditions. At day 30, LMNO fusion organoids show less caspase activation than neural organoids alone, suggesting that the meningeal tissue secretes factors that could prevent cell death during the static conditions of the LMNO fusion protocol and that the factors generated from mouse tissue affect human cells. The LMNO fusion system may benefit from experimentation with other aspects of the fusion model as well. For example, both the time post-differentiation of the organoid and the developmental stage of the mice can be manipulated to interrogate different questions. Fusing leptomeningeal tissue from an earlier time in development with a more developed neural organoid may be used to examine how signals from the developing brain impact leptomeningeal maturation. Further, fusion starting with an older neural organoid may allow for examination of interactions between human iPSC-derived glia and leptomeningeal tissue. Overall, the LMNO fusion model shows that human neural organoids can functionally integrate into mouse meningeal tissue and that this platform can be used to reveal the intricate meninges-brain signaling networks that underlie early brain development.

## Methods

### Animals

All mice were housed in specific-pathogen-free facilities approved by AALAC and were handled in accordance with protocols approved by the IACUC committee on animal research at the University of Colorado, Anschutz Medical Campus. The following mouse line was used in this study: Col1a1-GFP^17^ on a C57/BLK6J background. GFP+ embryos were identified by fluorescence detection using a stereoscope equipped with a fluorescent light source.

### Bioinformatic Analysis

Published human embryonic and fetal brain single nuclei data sets were generously provided by Dr. Aparna Shah^25^ and Dr. Alexandro Trevino^26^ as R objects (Dr. Shah) or were available as raw single cell data from the NeMO Repository at^20^ and GEO: GSE162170. From individual data sets containing nuclei isolated from Carnegie Stage (CS) 12-22 (correspond to gestational weeks 6-10) cortex/telencephalon, hindbrain, midbrain or thalamus, clusters with expression of putative meningeal mesenchymal cell enriched genes (*COL1A1, COL1A2, LUM, S100A6, CRABP2, TBX18, FOXC1, DCN, NNAT, ALCAM, and BGN)*, endothelial genes (*CLDN5*, *PECAM1*, *SOX17*), mural cell genes (*RGS5, ABCC9, KCNJ8*) or macrophages/monocytes/microglia (*MRC1, CD74, AIF1, P2YR12*) were subset, integrated and re-clustered. This dataset was further subset to remove a cluster of putative neural cells and reclustered at 0.8 resolution leaving clusters of putative mesenchymal (combined meningeal mesenchymal and pericyte markers), endothelial and macrophages/monocytes/microglia. From a data set containing nuclei isolated from GW16-22 human cortex, previously annotated clusters 16 (‘microglia’), 19 (‘pericyte’), 20 (‘endothelial cell)’, and 22 (‘vascular and leptomeningeal cell’) were subset and cluster analysis was performed. Analysis of putative meningeal mesenchyme, pericyte/mural cell, monocyte/macrophage/microglia and endothelial markers in the data set showed the presence of clusters with neural identity. This dataset was further subset to remove a cluster of putative neural cells and reclustered at 0.8 resolution leaving clusters of meningeal, pericyte/mural cell, monocyte/macrophage/microglia and endothelial cells.

### Human iPSCs

The WTC11 iPSC line used in this study was maintained in E8 medium in plates coated with Matrigel (Corning, cat # 354277) at 37°C with 5% CO2. Culture medium was changed daily. Cells were checked daily for differentiation and were passaged every 3-4 days using Gentle cell dissociation solution (STEMCELL Technologies, cat # 07174). All experiments were performed under the supervision of the Vanderbilt Institutional Human Pluripotent Cell Research Oversight (VIHPCRO) Committee. Cells were checked for contamination periodically.

### Neural organoids

Cerebral organoids were generated as previously described^3,4,16^ with some modifications. Briefly, organoids were generated using the STEMdiff™ Cerebral Organoid Kit (STEMCELL Technologies; Cat# 08571, 08570), supplemented with dual SMAD inhibitors. iPSCs were dissociated into single cells using Gentle Cell Dissociation Reagent (STEMCELL Technologies, cat # 07174) for 8 minutes at 37°C. Homogeneous and reproducible EBs were generated by using a 24-well plate AggreWell™ 800 (STEMCELL Technologies, cat # 34815). On Day 7, high-quality EBs were embedded in Matrigel (Corning, cat # 354277). On Day 10, the Matrigel coat was broken by pipetting up and down and the healthy organoids were transferred to a 60mm low attachment culture plate (Eppendorf, cat # 003070119). The plates were then moved to a 37°C incubator and to a Celltron benchtop shaker for CO2 incubators (Infors USA, cat # I69222) set at 85rpm. Full media changes were performed every 3–4 days for 15 days.

### Isolation of Leptomeninges

Embryos were collected from *Col1a1-GFP* pregnant dams at E16. Leptomeningeal tissue was dissected using previously established methods^14^; briefly, a single lateral cut is made from the back of the head towards the eye, and the skin and calvarium is lifted dorsally exposing the brain surface and GFP+ leptomeninges. Whole pieces of leptomeninges were then peeled off the forebrain (telencephalon) using a dissecting scope and then stored in ion-free HBSS on ice until fusion with organoids.

### Generation of LMNO fusions

Leptomeninges from mice at E16 of embryonic development were isolated (described above) and seeded together with neural organoids in a round-bottom tube placed within a modified 6-well plate. The leptomeninges fused to forebrain neural organoids within 6 days and were kept in culture for 30 to 60 days, with complete media changes and imaging using a fluorescent stereoscope every other day.

### Immunohistochemistry and RNAscope

At day 30 and 60, leptomeningeal-neural organoid fusions were collected and sectioned prior to immunohistochemistry and detection of mRNA using RNAScope. Briefly, LMNO fusions were fixed in 4% paraformaldehyde for 15 minutes at 4°C, washed in PBS, then incubated in a 20% sucrose gradient at 4°C prior to embedding and flash-freezing in a solution of 7.5% gelatin/10% sucrose. LMNO fusions were sectioned on a cryostat (Leica CM1950) at a thickness of 15μm. For detection of leptomeningeal components and assaying for cell death, the following antibodies were used at a dilution of 1:100: CD206 (goat, R&D Systems, AF2535), Lyve1 (rat, R&D Systems, MAB2125), E-cadherin (mouse, BD Biosciences, 610181), Claudin-11 (rabbit, Invitrogen, 36-4500), CRABP2 (mouse, Sigma-Aldrich, MAB5488), Cleaved Caspase-3 (rabbit, Cell Signaling Technology, 9661). For immunolabeling of Cajal-Retzius like cells the following antibodies were used at a dilution of 1:200: Reelin (mouse, Sigma-Aldrich, MAB5366), and Mitochondria (mouse, Abcam, ab92824). For detection of *Cxcl12* transcript, the RNAScope Manual Detection Kit v.2.0 from ACDBio (Cat. No. 323100) and probe against mouse *Cxcl12* (Cat. No. 422711-C3).

#### Image acquisition

Confocal images were acquired on a Nikon Ti2 inverted light microscope equipped with a Yokogawa CSU-X1 spinning disk head, with 488 nm, 561 nm, and 647 nm excitation LASERs, and Photometrics Prime 95B sCMOS camera.

## Acknowledgements

We would like to thank Stellan Riffle and Caroline Bodnya for providing technical support during organoid generation. This work was supported by 1R35 GM128915-01NIGMS (VG), 1RF1MH123971-01 (VG and JS), F99NS125829 (GLR), and F31NS125875 (HEJ). All SIM and spinning disk confocal microscopy imaging and image analysis were performed in part using the Vanderbilt Cell Imaging Shared Resource, which is supported by NIH grants 1S10OD012324-01 and 1S10OD021630-01.

The authors declare no competing financial interests.

